# Single-Cell Atlas of Dorsal Root Ganglion Remodeling After Neuroma-Forming Nerve Injury Reveals Intervention-Specific Glial and Immune Programs

**DOI:** 10.64898/2026.05.24.726282

**Authors:** Casey L. Stewart, Michael M. Morris, Alexandra E. Halevi, Amy M. Moore, Valeria Cavalli, Oshri Avraham

## Abstract

Peripheral nerve injury, whether caused by a cut, crush, or excessive stretch, can cause disordered regeneration and formation of a painful neuroma at the injury site. Regrowing axons may end in a swelling, hypersensitive nerve stump, producing pain along the affected nerve distribution that may outweigh sensory or motor deficits. Although neuromas are common, treatment outcomes are inconsistent and the mechanisms that initiate and sustain neuroma-associated pain remain poorly defined. Many surgical and non-surgical approaches (excision with transposition, capping, sclerosis, cryoablation) are used, yet comparative studies have not identified a superior technique, highlighting biological heterogeneity and knowledge gaps. A proximal nerve crush (axonotmetic injury) performed upstream of a neuroma or transection is proposed to reshape axonal growth and interrupt retrograde injury signaling, potentially shifting the system toward a more regenerative, pain-resolving state. However, how proximal crush remodels long-term programs in the dorsal root ganglion (DRG), and how these changes relate to neuroma-like outcomes, are unclear. Here, we build a single-cell atlas of DRG remodeling after neuroma-forming injury and compare it with two interventions- proximal nerve crush and nerve resection, using scRNA-seq. By resolving transcriptional responses across sensory neurons, satellite glial cells, Schwann cells, and macrophages, we identify intervention-specific glial and immune programs that distinguish permissive regeneration from persistent, pain-associated states and nominate pathways for mechanism-guided neuroma therapies.

## Introduction

Peripheral nerve injury, either caused by a cut, a crush, or an excessive stretch, can lead to the formation of a painful neuroma at the site of injury. Following traumatic injury, regenerating nerves may form a swelling at the severed end, and patients can experience intense pain along the nerve territory. In many cases, this pain is more debilitating than the accompanying loss of sensation or motor impairment, making symptomatic neuromas a major clinical driver of chronic neuropathic pain and disability. Despite this clinical burden, the molecular basis of symptomatic neuroma formation and the transition from acute injury to persistent pain remain incompletely understood.

Current management strategies reflect this mechanistic uncertainty. Approaches including nerve transposition into muscle, neuroma capping with synthetic or biologic materials, chemical sclerosis, electrocautery, fascicle ligation, and cryoablation have all been used with variable success (Vernadakis, Koch, and Mackinnon 2003). Surgical excision of the neuroma, with or without nerve transposition, is frequently pursued, and meta-analysis indicates that surgery can provide clinically meaningful pain relief in many cases. Importantly, comparative analyses have not identified a single superior surgical technique, highlighting both biological heterogeneity and gaps in mechanistic understanding (Poppler et al. 2018). One approach that has been used intermittently in clinical practice is a proximal crush (axonotmetic) intervention, described as a treatment modality for painful hand neuromas with favorable results in selected settings(Domeshek et al. 2017; Tupper and Booth 1976). Experimental work supports the idea that axonotmetic and neurotmetic injuries engage distinct long-term transcriptional programs in the dorsal root ganglion (DRG). In particular, targeted gene analyses have suggested that transection can produce sustained upregulation of neuropathic pain-associated programs (including c-Jun/AP-1 linked responses), whereas expression after crush injury may normalize over time, consistent with a more permissive regenerative state (Kenney and Kocsis 1998). Mechanistically, a proximal crush is hypothesized to reshape subsequent axonal growth by interrupting retrogradely transported injury signals and thereby altering the growth competence state of adult neurons (Smith and Skene 1997). While this framework is compelling, the field lacks a high-resolution, cell type-resolved view of how these injury contexts differentially remodel the DRG and how non-neuronal compartments contribute to persistent pain versus resolution. Sensory neurons mount regeneration-associated gene (RAG) programs that support axon regrowth (He and Jin 2016; Mahar and Cavalli 2018; Liu et al. 2011). However, peripheral nerve injury elicits a coordinated response across the DRG microenvironment, engaging glia, macrophages, and other non-neuronal cells (Avraham et al. 2020; Avraham et al. 2021; Jessen and Mirsky 2016; Zigrnond and Echevarria 2019). Neuroma formation and pain are therefore unlikely to be neuron-autonomous. Yet, most prior studies have relied on bulk tissue analyses or limited marker panels, leaving it unclear which cell states and pathways distinguish neuroma-forming injury from interventions proposed to mitigate neuroma pain.

In this study, we use single-cell RNA sequencing to define cell type-specific transcriptional programs in the DRG following two interventions: proximal nerve crush versus nerve resection following neuroma formation. We hypothesize that a proximal crush intervention modulates the maladaptive gene expression programs associated with painful neuroma-like states by promoting a pro-regenerative, pain-resolving environment. Across cell types, we observe a conserved core injury response, however proximal crush and neuroma resection elicit distinct glial state transitions and reshape intercellular signaling networks. Extracellular matrix (ECM) and adhesion pathways, particularly syndecan and integrin-centered modules, emerge as dominant features of neuron-satellite glial cell (SGC) communication, and functional evidence implicates Sdc4 as a key organizer of neuron-glia interactions.

By resolving responses across sensory neurons, satellite glia, Schwann cells, and macrophages, our work provides a comparative atlas of DRG remodeling and identifies molecular pathways that differentiate permissive versus persistent pain-associated injury states. Together, these data offer a mechanistic framework for symptomatic neuroma formation and highlight candidate targets for more effective, mechanism-guided therapies.

## Results

### Single-cell transcriptomic profiling of dorsal root ganglia across intervention approaches in a rat neuroma model

A visible neuroma formed at the distal end of the transected sciatic nerve approximately two weeks after injury (Fig. 1A). Histomorphometric analysis of the transected nerve stump at this timepoint revealed increased intrafascicular collagen deposition and disrupted axonal organization (reduced structural anisotropy) compared to uninjured sciatic nerve (Fig. 1B,C). These features are consistent with neuroma formation and resemble histopathologic changes reported in human neuromas (Egami et al. 2016; Vora et al. 2007; Yan et al. 2012). To define transcriptional responses in the DRG at single-cell resolution, and to compare proximal nerve crush with nerve resection following transection and neuroma formation, we harvested lumbar DRGs from adult male Lewis rats for single-cell RNA sequencing (scRNA-seq) (Fig. 1D). Across all conditions and replicates, we profiled 63,570 cells from two biological replicates per group: uninjured (15,899 cells), cut (9,731 cells), crush (19,323 cells), and resect (18,627 cells) (Fig. 1E). After quality filtering, we detected 25,267 genes in total (see filtering criteria in the methods). Unbiased graph-based clustering identified eight major annotated cell populations, plus an N/A cluster across control and injury conditions (Fig. 1F). Cluster identity was assigned using differential expression of cluster-enriched markers (ANOVA; fold-change threshold >1.5), revealing canonical DRG cell types: sensory neurons (Isl1, Tubb3, Gal, Tac, Prph), satellite glial cells (SGCs) (Fabp7, Cdh19), endothelial cells (Pecam1/Cd31, Flt1, Cldc5), Schwann cells (SC) myelinating and non-myelinating (Ncmap, S100b, Mag, Pmp22), pericytes (Kcnj8/Kir6.1, Pdgfrb), mesenchymal cells (Cldn11, Apod, Pdgfra), macrophages (Aif1/Iba1, Cd163, Mrc1), and connective tissue cells (Col1a1, Dcn). Visualization of all cells by UMAP showed that SGCs comprised the largest cluster across the dataset (Fig. 1F). Comparing cell-type composition across conditions revealed a consistent reduction in the proportion of SGCs in all three injury conditions, accompanied by an increased proportion of macrophages relative to uninjured controls (Fig. 1G,H).

**Figure 1.**
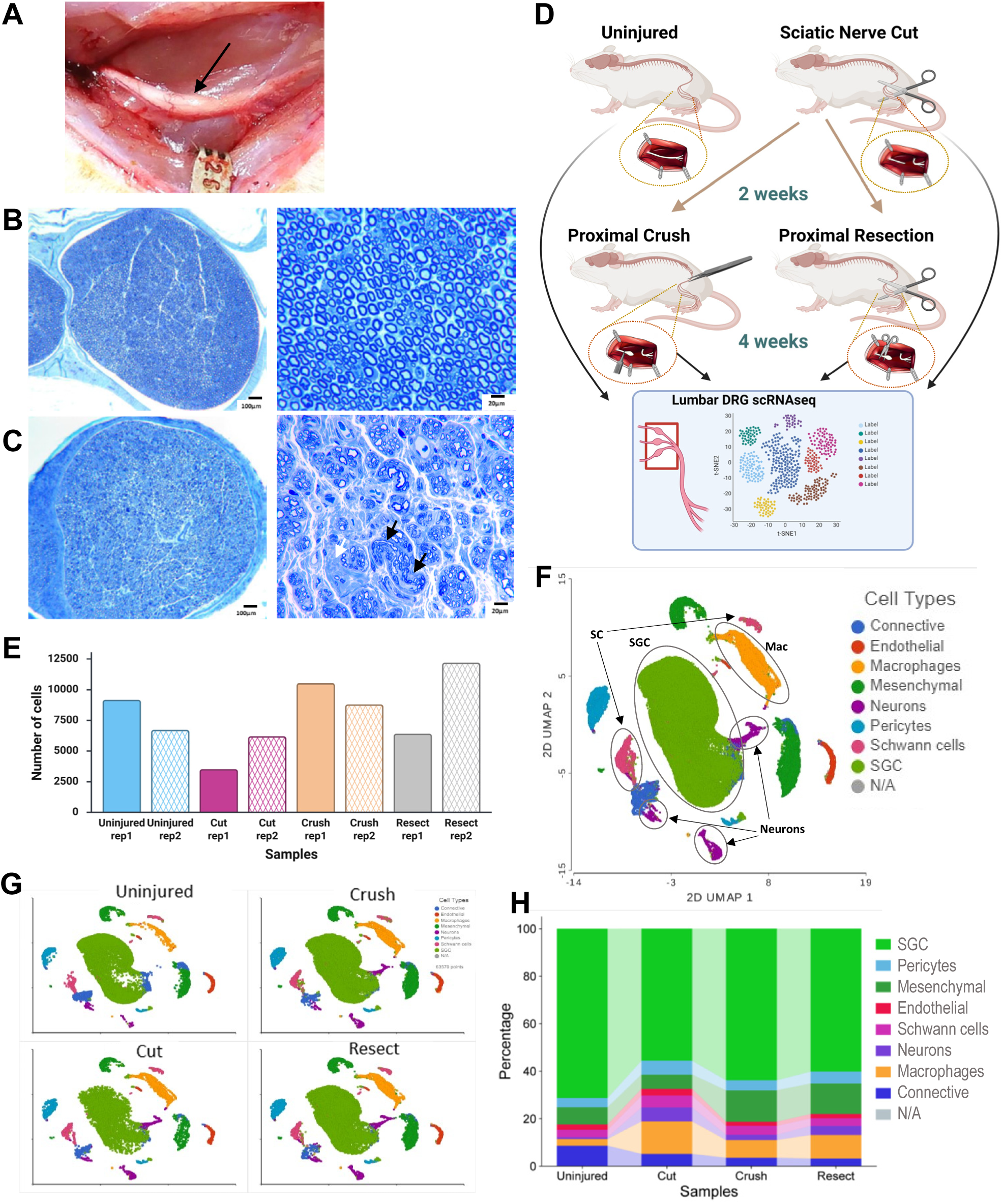
Neuroma rat model and intervention paradigm. **A**) Gross appearance of a terminal neuroma at the distal stump two weeks after sciatic nerve transection. **B**) Representative light microscopy images of uninjured rat sciatic nerve at low and high magnification, showing orderly fascicular architecture with uniformly distributed axons and relatively consistent axon caliber. **C**) Representative images of a terminal neuroma two weeks after sciatic nerve transection, demonstrating increased intrafascicular collagen deposition (white arrow) and disrupted axonal organization with reduced anisotropy (black arrows). Scale bars: 100 µm (low magnification) and 20 µm (high magnification). **D**) Experimental design and sampling workflow. After sciatic nerve transection and neuroma formation (cut), animals underwent one of two secondary interventions at two weeks: proximal sciatic nerve crush (crush) or neuroma resection (resect). L4-L5 DRGs were harvested from all injury groups and uninjured control and processed for scRNA-seq. **E)** Number of cells captured per treatment condition and batch. **F)** UMAP of all profiled DRG cells with major cell populations annotated based on canonical marker genes, identifying eight clusters (neurons, satellite glia, Schwann cells, macrophages, endothelial cells, pericytes, mesenchymal cells, connective tissue cells) **G)** UMAP visualization of clusters separated by treatment group (uninjured, cut, crush, resect). **H**) Cell-type composition across conditions shown as the fraction of each cluster within uninjured, cut, crush, and resect groups. n = 2 biological replicates per condition.

### Resection promotes metabolic and glial differentiation programs in SGCs, whereas proximal crush induces inflammatory signaling

SGCs form an ensheathing layer around sensory neuron somata, providing structural and metabolic support (Hanani and Spray 2020; Meriau, Kuruvilla, and Cavalli 2025). Following peripheral nerve injury, SGCs can influence neuronal excitability and pain signaling, and they participate in repair-associated programs, including induction of metabolic and supportive pathways (Andreeva et al. 2022; Avraham et al. 2020; Avraham et al. 2021; Jager et al. 2020). To define SGC responses two weeks after sciatic nerve transection, and to compare the effects of a subsequent proximal crush versus resection of the neuroma, we quantified differential gene expression within the SGC cluster (40,275 cells). Differentially expressed (DE) genes were identified relative to uninjured controls using an FDR ≤ 0.05 and fold change (FC) ≥2 (Supp. table 1). Following nerve cut alone, SGCs exhibited a robust transcriptional response, with 697 genes upregulated and only 1 gene downregulated. In contrast, the proximal crush condition showed a markedly smaller set of changes (59 upregulated genes), while neuroma resection induced 156 upregulated genes and 5 downregulated genes. Pathway enrichment analysis of the cut condition highlighted strong activation of metabolic programs, particularly pathways related to α-linolenic acid, cholesterol, and fatty acid metabolism, alongside signatures of glial differentiation (Fig. 2A). Adhesion-related pathways, including cadherins and adherens junction components, were also enriched, consistent with injury-associated remodeling of SGC organization within the DRG and increased metabolic support to injured neurons. The only gene that was significantly downregulated was Lumican (Lum), that encodes a small leucine-rich proteoglycan (SLRP) and known to play an important structural and regulatory role in the extracellular matrix. Strikingly, when a proximal crush was performed two weeks after transection, SGCs preferentially upregulated immune-related pathways, including chemokine and cytokine signaling (Fig. 2B). By comparison, resection of the neuroma primarily reinforced metabolic and trophic programs, with enrichment for pathways linked to metabolism, neurotrophin-related signaling, and SGC proliferation/differentiation (Fig. 2C). Consistent with these pathway-level patterns, overlap analysis showed substantial similarity between the cut and resection responses (85 shared genes), but minimal overlap between cut and crush (6 shared genes) (Fig. 2D). A heatmap of SGC DE genes further supported this divergence (Fig. 2E). Glial differentiation-associated genes (Sox10, Notch1, Ednrb, Sox2) and metabolic genes (Fabp7, Fabp5, Fasn) were induced after cut but not after proximal crush. Conversely, inflammatory mediators (Cxcl2, Ccl4, Cxcl13) were strongly upregulated after proximal crush but showed little to no induction after initial cut. These results suggest that neuroma resection predominantly engages metabolic and differentiation programs in SGCs, whereas proximal crush shifts SGCs toward an inflammatory transcriptional state. Given that cytokine secretion and inflammatory signaling can contribute to peripheral nerve repair and regeneration, these SGC responses may represent distinct, intervention-dependent mechanisms shaping recovery after injury (Zhang et al. 2025; Gaudet, Popovich, and Ramer 2011). To directly assess how post-neuroma interventions influence molecular changes relative to the cut condition, we compared proximal crush and resection to cut injury. In SGCs, both interventions resulted in relatively few additional transcriptional changes beyond those induced by the cut. Specifically, three genes were upregulated following crush and five following resection, with only one shared gene, Lumican (Lum), which is involved in extracellular matrix (ECM) organization. Notably, Lum was also the only gene significantly downregulated in the crush versus uninjured condition (Supp. Fig. 1A, Supp. table 1). Crush selectively induced immune-related genes, including Ccl4 and Cxcl2, consistent with enhanced chemokine signaling. In contrast, resection preferentially upregulated genes associated with regenerative programs, including Epcam (stem cell differentiation) and Bcas1 (myelination), suggesting activation of repair-associated processes. Analysis of downregulated genes revealed 21 genes in crush and 13 in resection, with six shared genes. Pathway analysis indicated that both interventions suppressed gene programs associated with cell-cell and ECM interactions relative to the cut condition (Supp. Fig. 1B), consistent with remodeling of neuron-SGC coupling following intervention.

**Figure 2.**
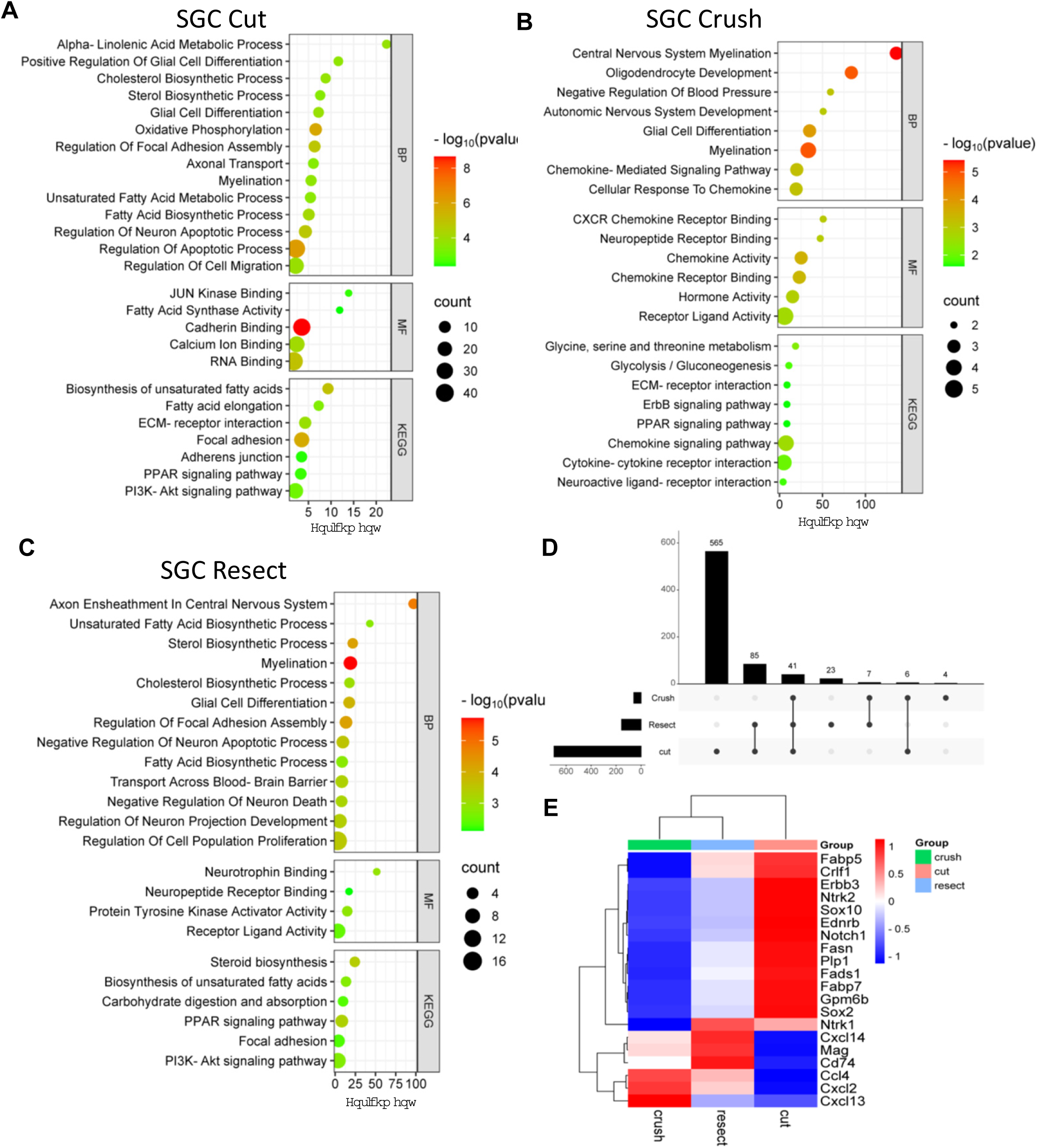
Neuroma interventions differentially reprogram satellite glial transcriptional programs. **A–C**) Pathway enrichment analysis of differentially expressed genes in the SGC cluster two weeks after injury, shown for **A**) nerve cut, **B**) proximal crush, and **C**) neuroma resection, each compared to uninjured controls. **D**) UpSet plot showing overlap of upregulated genes across the three intervention groups relative to uninjured controls, highlighting shared and condition-specific SGC responses. **E**) Heatmap of selected SGC differentially expressed genes representing key enriched pathways, illustrating intervention-dependent transcriptional signatures across cut, crush, and resection conditions.

### DRG Schwann cells upregulate cholesterol biosynthesis programs after nerve injury

Schwann cells (SCs) are central drivers of peripheral nerve repair. After injury, they dedifferentiate and activate repair programs that support debris clearance, guide axon regrowth, and ultimately enable remyelination (Jessen and Mirsky 2016). Although SCs primarily reside along peripheral nerves, a subset is captured during DRG dissections because SCs ensheath the proximal axonal segments within DRGs. In our dataset, we identified 2,329 SCs (myelinating and non-myelinating), representing ∼4% of all profiled DRG cells. Differential expression analysis revealed a strong and consistent injury response in DRG-associated SCs across all intervention groups. After nerve cut, SCs upregulated 962 genes, with enrichment for metabolic pathways and glial differentiation, and downregulated 162 genes enriched for glutamine transport as well as p53- and HIF-1-associated signaling (Fig. 3A,B, Supp. table 1). Following proximal crush, SCs upregulated 1,248 genes enriched for metabolic programs, myelination, and differentiation, while 93 genes were downregulated, including gene sets associated with ion transport, p53 signaling, and cell adhesion (Fig. 3A,C). Neuroma resection similarly induced broad transcriptional remodeling, with 1,038 genes upregulated and enrichment for metabolic pathways and cellular organization programs (including adherens junction and cadherin-binding pathways), alongside 93 downregulated genes enriched for chemotaxis and transport-related functions (Fig. 3D). Notably, 357 genes were upregulated across all three injury conditions (Fig. 3A). Pathway enrichment of this shared response was dominated by lipid metabolism, with particularly strong representation of steroid and cholesterol biosynthesis genes encoding the proteins SQLE, EBP, NSDHL, SC5D, MSMO1, LSS, CYP51, TM7SF2, and FDFT1 (Fig. 3E). Although Schwann cells in the DRG are far away from the injury site, this expression signature is biologically consistent with the critical role of cholesterol in myelin membrane synthesis, as well as the broader importance of steroid and lipid signaling in repair Schwann cell-neuron interactions during remyelination (Rodriguez-Waitkus et al. 2003; Saher et al. 2005). In contrast to the conserved cholesterol biosynthesis response, several canonical myelination and Schwann cell differentiation markers showed injury-specific regulation. Myelination and Schwann cell differentiation-associated genes (S100b, Sox10, Sox2, Plp1) were robustly induced after nerve transection, but showed little or no comparable upregulation after proximal crush or neuroma resection (Fig. 3F). This suggests that, although these conditions share a core metabolic/cholesterol response, distinct intervention strategies drive divergent Schwann cell state trajectories. In contrast, adhesion and cell-cell organization genes (Nrxn2, Cdh1) were suppressed after the initial nerve cut but increased after proximal crush and resection, consistent with a shift toward tissue organization/remodeling rather than differentiation. To further resolve how interventions reshape Schwann cell responses relative to the neuroma-forming cut injury, we directly compared crush and resection to the cut condition. This analysis identified 278 upregulated genes in crush and 314 in resection, with 78 genes shared between the two interventions (Supp. Fig. 1C). These shared genes were enriched for pathways related to cellular organization, including cytoskeletal remodeling and vesicle-mediated processes, suggesting convergence on structural reorganization programs following intervention (Supp. Fig. 1D). Downregulated gene analysis revealed 121 and 129 genes in the crush and resection conditions, respectively, with 52 genes shared. Notably, crush injury was associated with upregulation of inflammatory pathways, including IL-17 and chemokine signaling, whereas these same pathways were downregulated following resection. Both interventions also showed reduced expression of genes involved in ion transport, particularly calcium and sodium transporter activity (Supp. Fig. 1C,D), indicating a shift away from electrophysiological signaling toward structural and metabolic remodeling.

**Figure 3.**
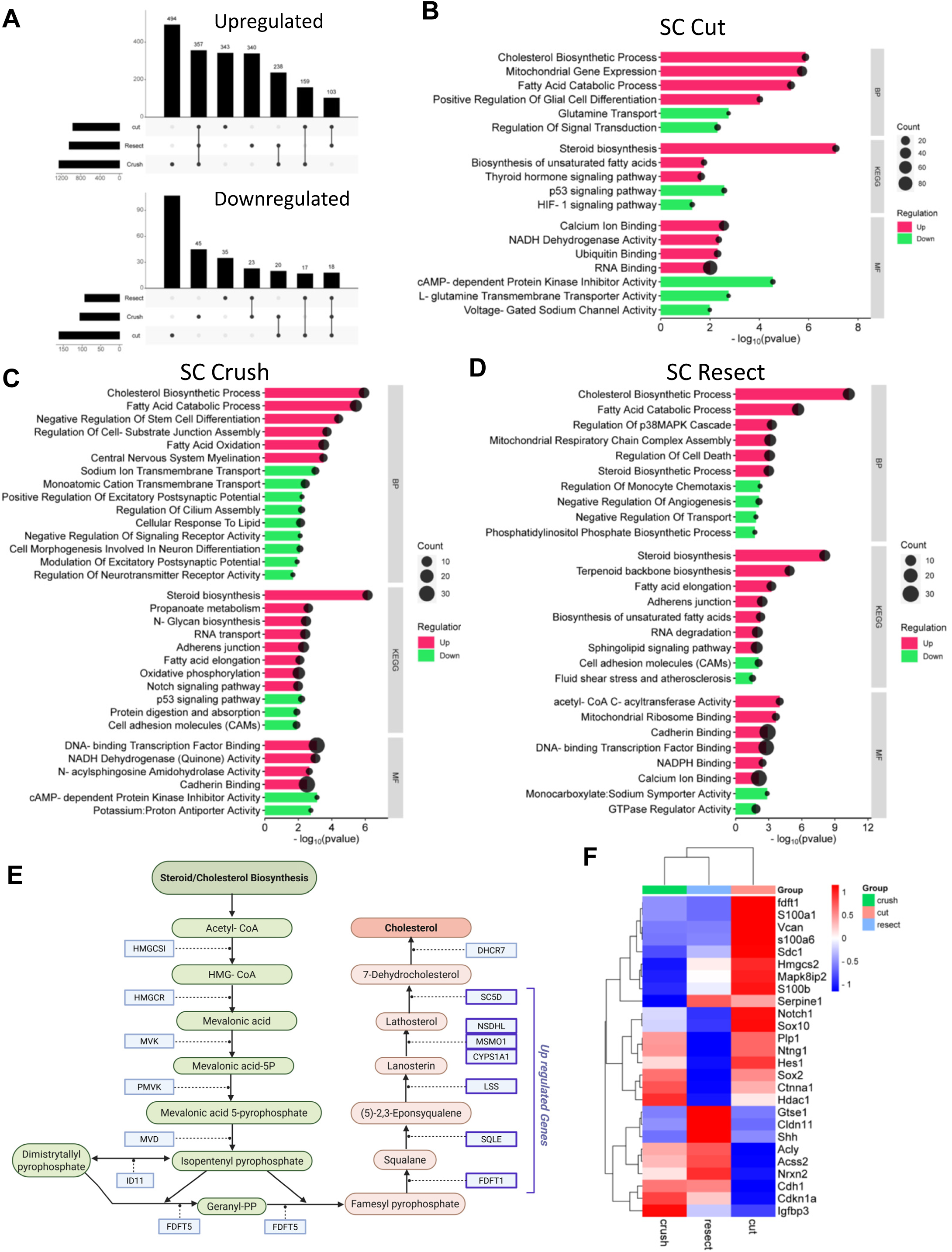
Schwann cell metabolic remodeling after nerve injury is centered on steroid/cholesterol pathways. **A**) UpSet plot showing overlap of differentially expressed genes in the SC cluster across injury conditions (cut, proximal crush, and neuroma resection), each compared to uninjured controls, highlighting shared and condition-specific responses. **B–D**) Pathway enrichment analysis of SC differentially expressed genes two weeks after injury, shown for **B**) nerve cut, **C**) proximal crush, and **D**) neuroma resection relative to uninjured controls. **E**) Steroid/cholesterol biosynthesis pathway map. Genes commonly upregulated across all three injury paradigms are highlighted in dark blue. F) Heatmap of selected SC genes from key enriched pathways, illustrating shared and injury-specific transcriptional changes across conditions.

### Neurons reorganize their structure after nerve damage

Sensory neurons are among the largest cells in the body. Their soma can range from ∼20 µm in nociceptors to ∼100 µm in proprioceptors. Because of this size, and their fragility, many neurons are easily damaged during standard whole-cell dissociation for scRNA-seq using microfluidics, which can lead to significant neuronal loss and biased sampling. Using isolated nuclei from DRGs substantially improves neuronal capture in single-cell assays (Renthal et al. 2020; Bhuiyan 2024). In our dataset, whole-cell scRNA-seq yielded 1,825 sensory neurons. We then performed differential expression analysis following the three injury paradigms. Each injury induced a large transcriptional response in neurons. After nerve cut, we detected 1,600 upregulated genes and 941 downregulated genes. A proximal crush produced an even larger induction, with 1,991 genes upregulated and 448 downregulated. Resection upregulated 1,633 genes, with 405 genes downregulated (Fig. 4A,B, Supp. table 1). While the glial cell response varied depending on the type of injury, neuronal responses shared a strong common core across all three injury conditions, despite differences in magnitude. A majority of the induced genes (1,022) were shared across all three injury conditions (Fig. 4A), suggesting that sensory neurons engage a conserved “injury program” regardless of the precise type of damage. Functional enrichment of these shared upregulated genes highlighted pathways consistent with neurons reorganizing their physical state, particularly cell migration, structural/cytoskeletal organization, and cell adhesion, including tight junction and adherens junction-related pathways (Fig. 4C-E). Interestingly, we also observed induction of gene sets associated with maintenance and remodeling of barrier properties, consistent with reorganization of the DRG blood-nerve barrier after injury. This included upregulation of canonical junction-associated genes such as Tjp1, Tjp2, Jam2, and Jam3 (Fig. 4F, Supp. table 1), supported by enrichment analysis (Evangelista et al. 2023). In contrast, many of the genes downregulated across injuries were enriched for ion transport pathways, particularly channels involved in calcium, sodium, and potassium handling (Fig. 4C-E; Supp. table 1). This pattern is consistent with neurons temporarily shifting away from an excitable, signaling-focused state and toward a remodeling/repair-associated state following damage (Renthal et al. 2020; Jang and Garraway 2024). In those studies, neuronal responses were examined 3–7 days after injury, and found that somatosensory neuronal subtypes mounted a broadly shared transcriptional program that both promoted axon regeneration and suppressed aspects of mature cell identity. Notably, this reprogramming was specific to neurons and was not observed in non-neuronal cells. It resolved over the course of approximately 7 days, coincident with target reinnervation and restoration of the original neuronal identity (Renthal et al. 2020). In contrast, our paradigm involved analysis at later time points, 2 and 4 weeks after injury, where the observed differences likely reflect more sustained functional adaptations rather than the acute transcriptional reprogramming described at earlier stages. Direct comparison of crush and resection to the cut condition revealed substantial additional transcriptional remodeling in neurons. Crush induced 746 upregulated genes and resection induced 899, with 275 shared genes enriched for ion transport pathways (Supp. Fig. 2A,B; Supp. table 1). Resection also resulted in a greater number of downregulated genes compared to crush (232 vs. 156), with 62 shared genes. Both interventions were enriched for pathways related to extracellular matrix organization and growth factor activity, suggesting enhanced structural remodeling and trophic signaling relative to cut injury. Interestingly, crush uniquely led to downregulation of inflammatory pathways, including interleukin-1, TNF, and IL-17 signaling, whereas these pathways were not significantly altered in the resection condition. Together, these results indicate that, while neurons maintain a conserved injury response, secondary interventions further modulate inflammatory and structural programs in an intervention-specific manner.

**Figure 4.**
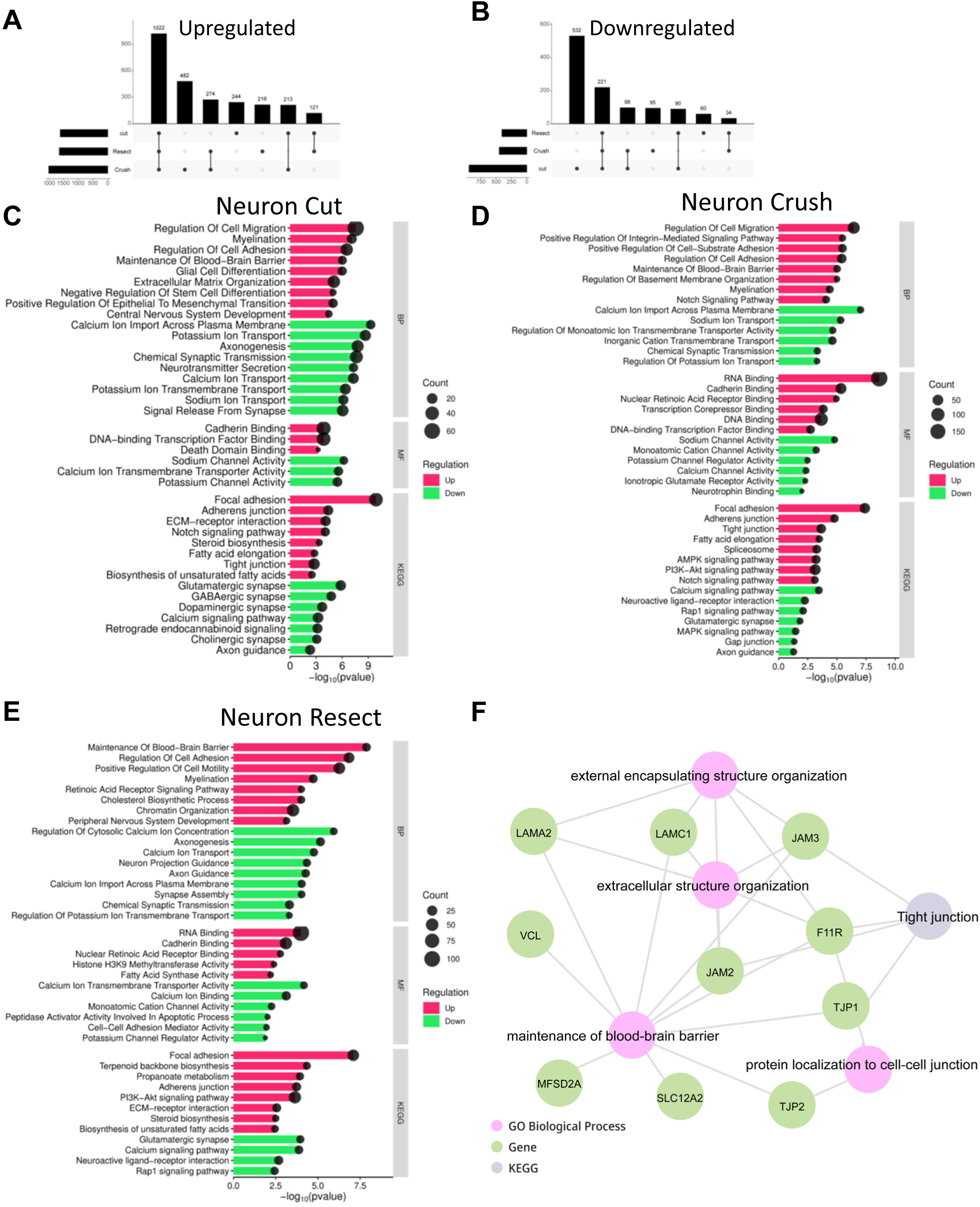
DRG neurons activate a conserved injury transcriptional program across interventions. **A-B)** UpSet plot showing overlap of up (A) and down (B) regulated genes in the neuronal cluster across injury conditions (cut, proximal crush, and neuroma resection), each compared to uninjured controls, highlighting shared and condition-specific responses. **C–E**) Pathway enrichment analysis of neuronal differentially expressed genes two weeks after injury, shown for **C**) nerve cut, **D**) proximal crush, and **E**) neuroma resection relative to uninjured controls. **F)** Enrichment analysis of the genes commonly regulated across all three injury paradigms in DRG neurons, highlighting shared biological programs induced by nerve injury.

### Injury-Evoked Macrophage Responses in the DRG

Resident macrophages comprise ∼3% of cells in uninjured DRGs, but their abundance increases markedly after nerve injury through a combination of local proliferation and immune cell infiltration. In our dataset, macrophage numbers rose approximately 3-4 fold following injury (Fig. 1H), consistent with prior reports in related nerve injury models(Avraham et al. 2021). Macrophages are known to support nerve repair at the injury site and to shape pain-related signaling within the DRG (Kwon et al. 2013; Niemi et al. 2013; Feng et al. 2023). Across all conditions, we sequenced 5,057 macrophages. Differential expression analysis revealed robust transcriptional remodeling after each injury type. Following nerve cut, macrophages upregulated 1,039 genes and downregulated 310 genes. A proximal crush induced 885 upregulated and 185 downregulated genes, while resection altered 748 genes, including 219 downregulated (Fig. 5A,B, Supp. table 1). Across all injuries, shared upregulated programs were strongly enriched for fatty acid oxidation and oxidative phosphorylation, with prominent induction of mitochondrial and electron transport chain components, including NDUFA subunits, SDH subunits, and ATP synthase subunits (Fig. 5C-F). Notably, resection uniquely triggered enrichment of gene sets associated with neuronal development (Fig. 5E), suggesting an injury-specific macrophage state that may be particularly engaged in neural remodeling cues. In contrast, downregulated pathways across injuries were dominated by inflammatory response signatures, including cytokine/chemokine signaling as well as TNF and IL-17 pathway-associated genes. One additional distinction emerged after proximal crush, which selectively showed downregulation of cell proliferation pathways and a lower macrophage fraction in the DRG (∼7%), roughly half of what we observed after nerve cut (Fig. 1H). Together, these data indicate that DRG macrophages expand after injury and undergo a shared metabolic shift, while specific injury types, particularly resection and proximal crush, drive distinct transcriptional features. To determine how interventions reshape macrophage responses relative to cut injury, we directly compared crush and resection conditions to the cut state. This analysis identified 81 upregulated genes in crush and 98 in resection, with 19 genes shared between the two conditions (Supp. Fig. 2C; Supp. table 1). In both conditions, macrophages upregulated genes associated with negative regulation of angiogenesis and modulation of inflammatory pathways, including interferon, interleukin, and broader cytokine signaling. Analysis of downregulated genes revealed 91 genes in crush and 103 in resection, with 44 shared genes (Supp. Fig. 2C,D). These findings suggest that, relative to cut injury, both interventions shift macrophages toward a more regulated inflammatory and tissue remodeling state.

**Figure 5.**
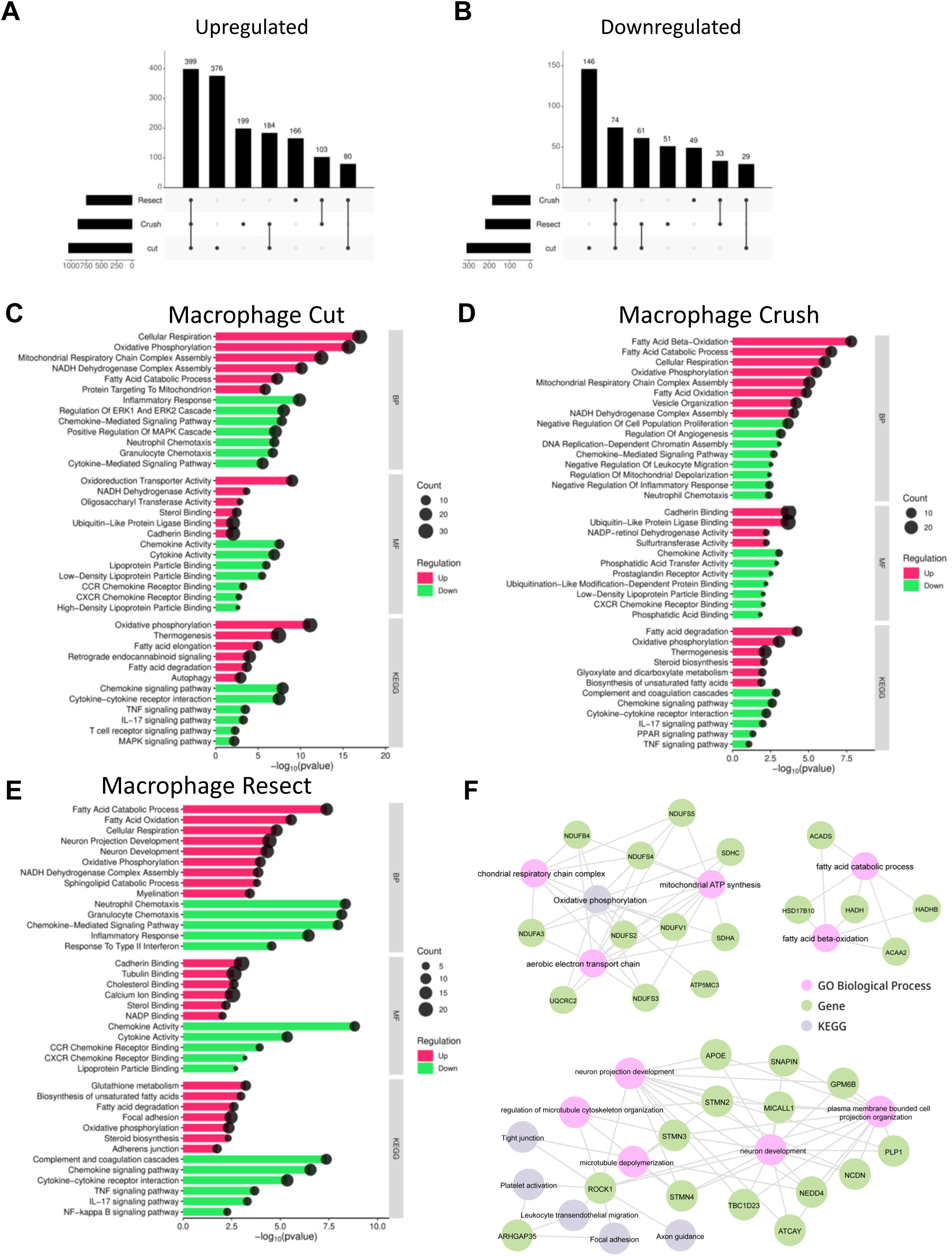
DRG macrophages undergo metabolic reprogramming and reduced inflammatory signaling after injury. **A-B)** UpSet plot showing overlap of up (A) and down (B) regulated genes in the macrophage cluster across injury conditions (cut, proximal crush, and neuroma resection), each compared to uninjured controls, highlighting shared and condition-specific responses. **C-E**) Pathway enrichment analysis of macrophage differentially expressed genes two weeks after injury, shown for **C**) nerve cut, **D**) proximal crush, and **E**) neuroma resection relative to uninjured controls. **F**) Enrichment analysis of genes commonly regulated across all three injury paradigms in DRG macrophages, highlighting shared biological programs.

### Ligand-Receptor Network Analysis Reveals Dynamic DRG Communication Programs After Injury

We next asked how nerve injury reshapes signaling between sensory neurons, glial populations (SGCs and SCs), and macrophages within the DRG. To infer intercellular communication, we performed ligand-receptor network analysis using scRNA-seq expression profiles, an approach that enables systematic characterization of condition-specific signaling relationships in vivo (Nagai et al. 2021; Moratalla-Navarro, Moreno, and Sanz-Pamplona 2023). Two weeks after a nerve cut, the dominant signaling changes were concentrated within and between neurons and SGCs, with 94 upregulated ligand-receptor pairs relative to uninjured controls (Fig. 6A-C, supp. table 2). In contrast, proximal crush shifted the interaction landscape toward neuron-macrophage communication, which accounted for ∼45% of upregulated interactions (37 ligand-receptor pairs), alongside substantial neuron-Schwann cell interactions (29 pairs) (Fig. 6A-C, Supp. table 2). Following resection of the neuroma, we observed a similarly prominent immune and glial component, with 42 neuron-macrophage and 44 neuron-SC upregulated interactions (Fig. 6A-C). Despite these injury-specific patterns, the response contained a shared core, where across all three injury conditions, we identified 20 common upregulated ligands and 14 common upregulated receptors (Fig. 6D,E, Supp. table 2). This suggests a conserved DRG “injury communication module” that is deployed broadly, with each injury paradigm layering on distinct interaction programs. We then focused on the highest-confidence pathways mediating communication between neurons and SGCs, as well as within these populations.

**Figure 6.**
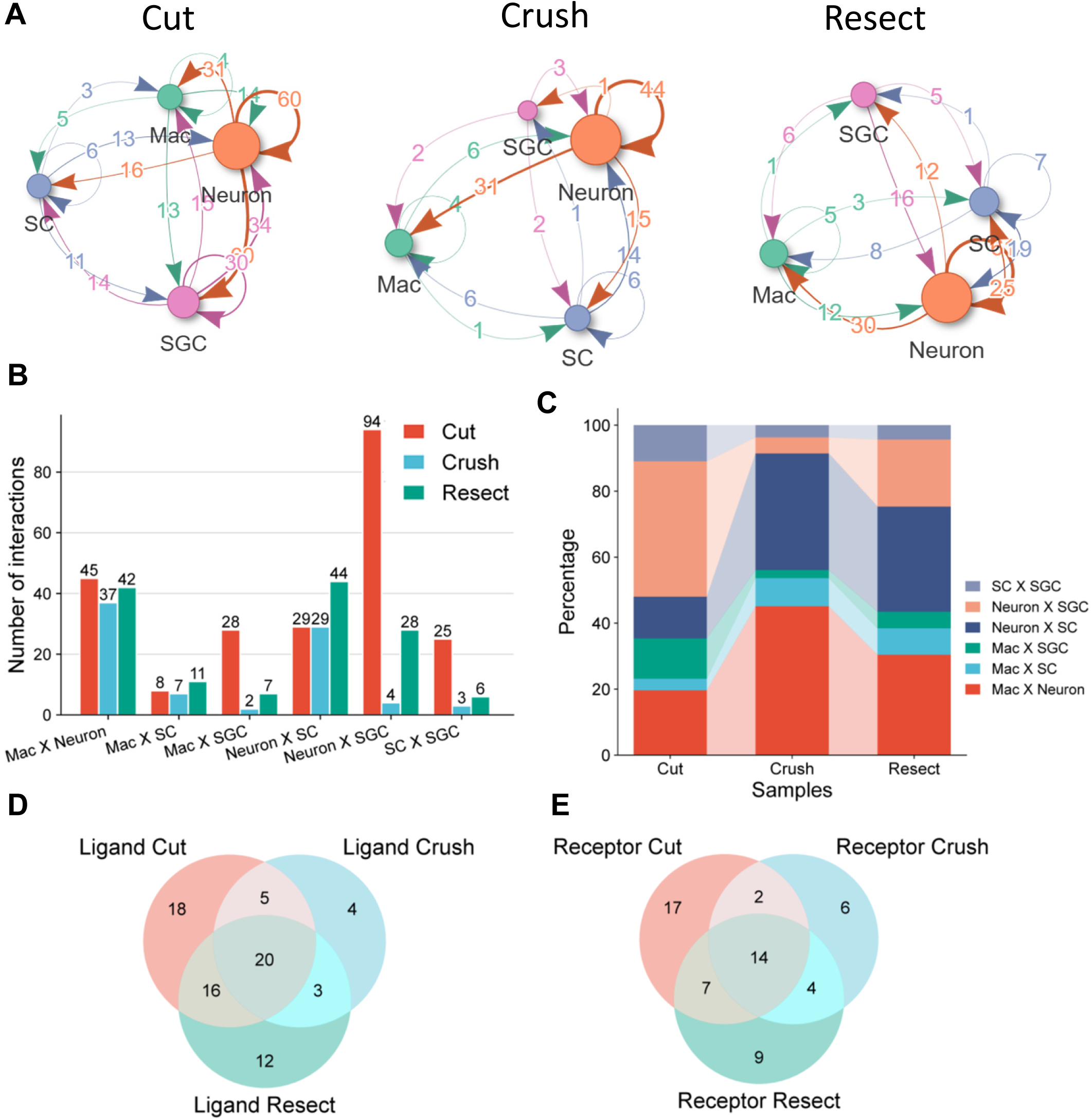
Ligand-receptor analysis reveals dynamic neuron-glia-immune signaling after injury. **A**) Ligand-receptor network inference across major DRG cell populations (neurons, SGCs, SCs, and macrophages) using scRNA-seq expression profiles from each injury condition. **B**) Quantification of significantly upregulated ligand–receptor interaction pairs identified in each condition relative to uninjured controls. **C**) Composition of interactions, showing the fraction of interactions occurring between each sender- receiver cluster combination across conditions. **D-E**) Venn diagram comparing upregulated **D**) ligand and **E**) receptor genes across injury paradigms.

### ECM–Syndecan/Integrin Networks Dominate Neuron-SGC Interactions After Nerve Injury

We next examined how sensory neurons and SGCs communicate across injury paradigms, focusing on upregulated ligand-receptor (LR) pairs inferred from scRNA-seq data. After nerve cut, neuron-SGC communication was the most prominent interaction class. We identified 94 upregulated neuron-SGC LR pairs, alongside 60 neuron-neuron and 30 SGC-SGC interactions (Fig. 7A). Among the most recurrent receptors were members of the syndecan family (Sdc2, Sdc3, Sdc4), a group of transmembrane heparan sulfate proteoglycans that function as co-receptors and modulate cell adhesion, migration, and signal transduction. We also observed strong representation of integrin receptors (Itgb1, Itgb4, Itga7), heterodimeric extracellular matrix (ECM) receptors that act as key sensors for adhesion, migration, and downstream signaling. Consistent with this receptor landscape, many of the top ligands were ECM-associated, including collagens (Col1a1, Col4a1, Col5a1, Col11a1) and laminins (Lamb2, Lamb1, Lama4, Lamc1), which provide structural support and regulate cell anchoring, migration, and tissue remodeling during repair (Fig. 7A). Pathway-level analysis of neuron-SGC signaling after nerve cut was enriched for ECM organization, receptor tyrosine kinase signaling, and syndecan/laminin-related interactions (Fig. 7B; Supp. table 3). Notably, Sdc4 and Itgb1 emerged as central receptors expressed in both neurons and SGCs, consistent with the idea that they serve as hubs integrating multiple ECM-derived cues. In contrast, after proximal crush, neuron-SGC signaling was markedly diminished, with only 4 upregulated neuron-SGC interactions, while neuron-neuron interactions expanded to 44 upregulated pairs (Fig. 7C, Supp. table 3). This shift suggests a relative disengagement of neuron-SGC coupling and a stronger emphasis on signaling within the neuronal compartment. The remaining upregulated programs still largely mapped to ECM-related pathways, consistent with ongoing structural remodeling (Fig. 7D). After resection, neuron-SGC communication demonstrated more neuron-SGC communication relative to crush, with 29 upregulated pairs and 61 neuron-neuron interactions (Fig. 7E, Supp. table 3). As with nerve cut, resection-associated signaling was enriched for syndecan and integrin receptors and their ECM ligands (collagens and laminins), indicating reactivation of adhesive ECM-receptor communication modules. Strikingly, resection was the only injury condition that robustly induced pathways linked to direct cell-cell communication and cell junction organization, involving ligands such as Cdh1 and F11r together with integrin-associated receptors (Fig. 7F). This pattern suggests that resection may uniquely promote a cell-cell contact and junctional signaling program that could be relevant for regeneration-associated tissue reorganization.

**Figure 7.**
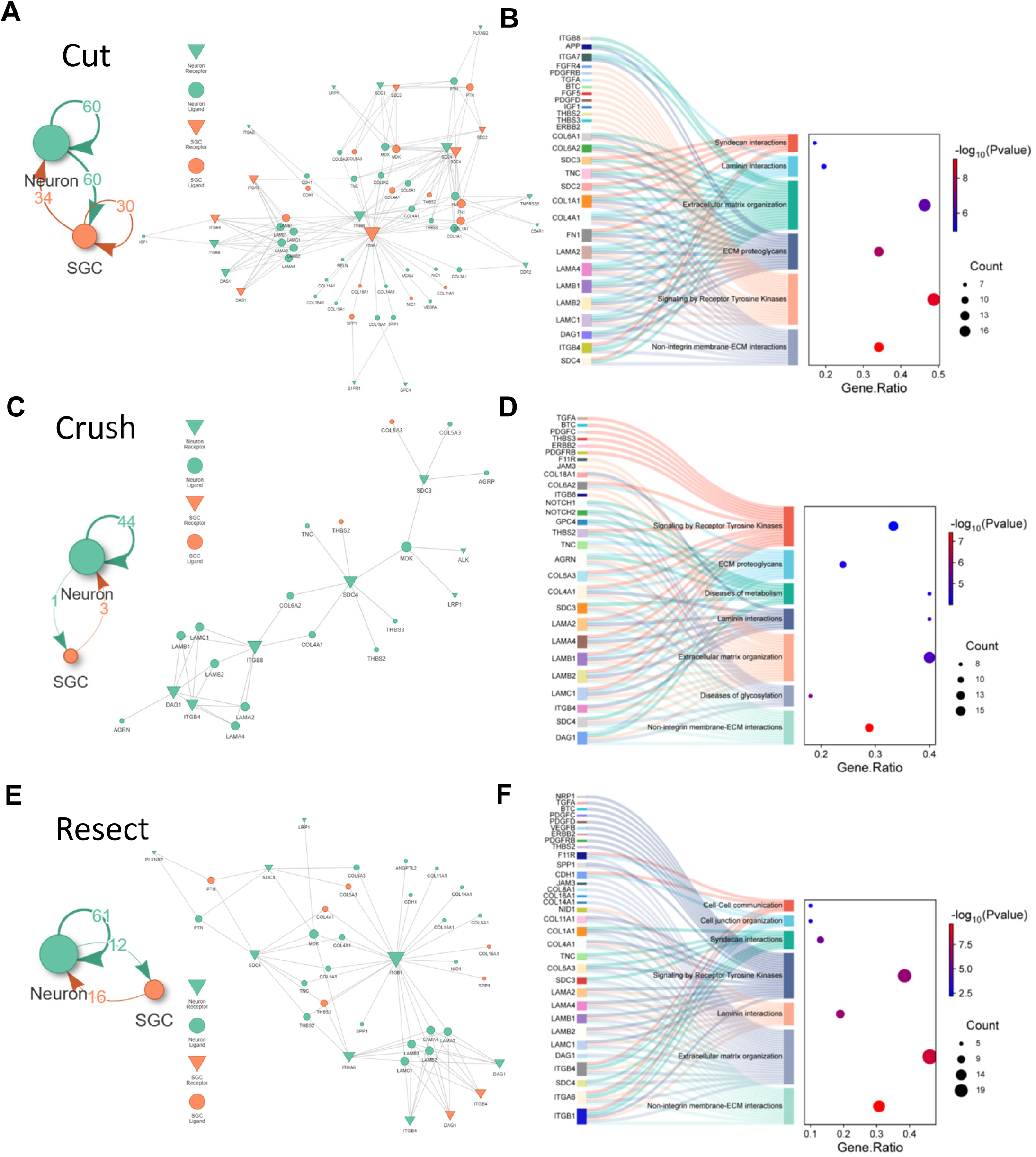
Injury-specific remodeling of neuron-SGC ligand-receptor networks. **A**) Ligand-receptor interaction network between DRG neurons and SGCs inferred from scRNA-seq data after nerve cut. Neuron-associated nodes are shown in teal and SGC-associated nodes in orange. Node size is proportional to degree (number of connections). Inverted triangles indicate membrane receptors, and rounded nodes indicate secreted ligands. **B**) Pathway enrichment summary for neuron-SGC ligand-receptor pairs after nerve cut, displayed as a Sankey plot linking enriched pathways to contributing genes, alongside a dot plot summarizing pathway significance and gene counts (dot size reflects the number of genes per pathway, color indicates enrichment p-value). **C**) Neuron-SGC ligand-receptor network after proximal crush **D**) Pathway enrichment and gene-level contributors for neuron-SGC ligand-receptor pairs after proximal crush. **E**) Neuron-SGC ligand-receptor network after neuroma resection. **F**) Pathway enrichment and gene-level contributors for neuron-SGC ligand-receptor pairs after neuroma resection.

### Adhesion Molecules Stabilize Neuron-SGC Interactions in a DRG Spot Co-culture Model

Because our transcriptional and ligand-receptor analyses identified ECM-receptor signaling as a dominant injury-regulated communication axis between neurons and SGCs, we next asked whether candidate adhesion receptors emerging from this analysis are functionally required for neuron-SGC organization. Among these candidates, Sdc4 stood out because it was expressed in both neurons and SGCs and was connected to multiple injury-induced ECM ligands, including collagens and thrombospondins. We therefore used shRNA-mediated knockdown in DRG spot cultures to test whether Sdc4 contributes to the spatial organization of SGCs around neuronal clusters. In the spot co-culture DRG model, dissociated DRG cells from E13.5 mouse embryos are plated into 24-well plates and seeded as a dense “spot” in the center of each well (Avraham et al. 2020; Avraham et al. 2021). In this system, neurons and glia initially distribute randomly, but over time they self-organize into a stereotyped architecture that is well-suited for quantifying neuron-glia adhesion and spatial patterning. DRG cells were transduced with viral particles encoding shRNAs targeting candidate adhesion genes, and cultures were analyzed five days later for changes in cell localization and organization (Fig. 8A). At DIV1 (days in-vitro), neurons and SGCs were intermingled within the spot. By DIV5, neurons formed prominent clusters, and SGCs wrapped neuronal somata within the spot while largely avoiding axons extending beyond the spot boundary (Fig. 8B). This transition provides a clear, quantifiable readout of neuron-SGC adhesion and positioning. We focused on three candidates with strong expression and/or injury regulation: Syndecan-4 (Sdc4), a key receptor repeatedly identified across injury paradigms in our ligand-receptor analysis (Fig. 7) and highly expressed in both sensory neurons and SGCs (Supp. table 1). The other candidates were Secreted phosphoprotein 1 (Spp1/osteopontin), a multifunctional ECM-associated glycoprotein implicated in ECM remodeling, adhesion, and migration, highly expressed in sensory neurons (Ewan et al. 2021) and upregulated after resection (Supp. table 1) and Amyloid precursor protein (App), a well-established neuronal protein important for developmental organization and plasticity, and expressed during DRG development (Avraham et al. 2022) and upregulated after nerve cut in SGCs (Fig. 7; Sup. table 1). To assess SGC localization, we stained cultures for the SGC marker FABP7. In control cultures, FABP7+ cells were largely restricted to the spot region, consistent with the expected organization. In contrast, Sdc4 knockdown produced a striking redistribution of SGCs, with FABP7+ cells appearing outside the spot boundary (Fig. 8C). Quantification confirmed a significant increase in FABP7+ cells outside the spot in cultures treated with Sdc4 shRNA and App shRNA, whereas Spp1 knockdown did not significantly differ from control (Fig. 8D, Supp. table 4). Together, these results support a model in which Sdc4, and potentially App -centered ECM signaling is not only associated with injury states but functionally contributes to neuron-SGC adhesion and organization. To connect these functional effects to our ligand-receptor networks, we next examined potential Sdc4 ligands expressed in DRG cell populations. We identified 10 candidate ligands, including multiple ECM components, such as collagens (Col1a1, Col4a1, Col6a1, Col6a2) and thrombospondins (Thbs2, Thbs4) (Supp. Fig. 3). Notably, Thbs4 was upregulated in SGCs, SCs, and macrophages across injury models, whereas Thbs2 showed induction primarily in SGCs and neurons. Additional candidate ligands included Pleiotrophin (Ptn), which was induced in SGCs, SCs, and neurons. Collagen transcripts were strongly induced in neurons and were most prominent in SGCs after cut injury. Overall, these perturbation experiments and ligand expression patterns suggest that ECM-receptor interactions (including Sdc4-centered signaling) are not merely correlated with injury states but can causally influence neuron-SGC organization. This supports the idea that adhesion and communication programs activated after peripheral nerve damage may contribute to the coordinated cellular remodeling required for repair and regeneration.

**Figure 8.**
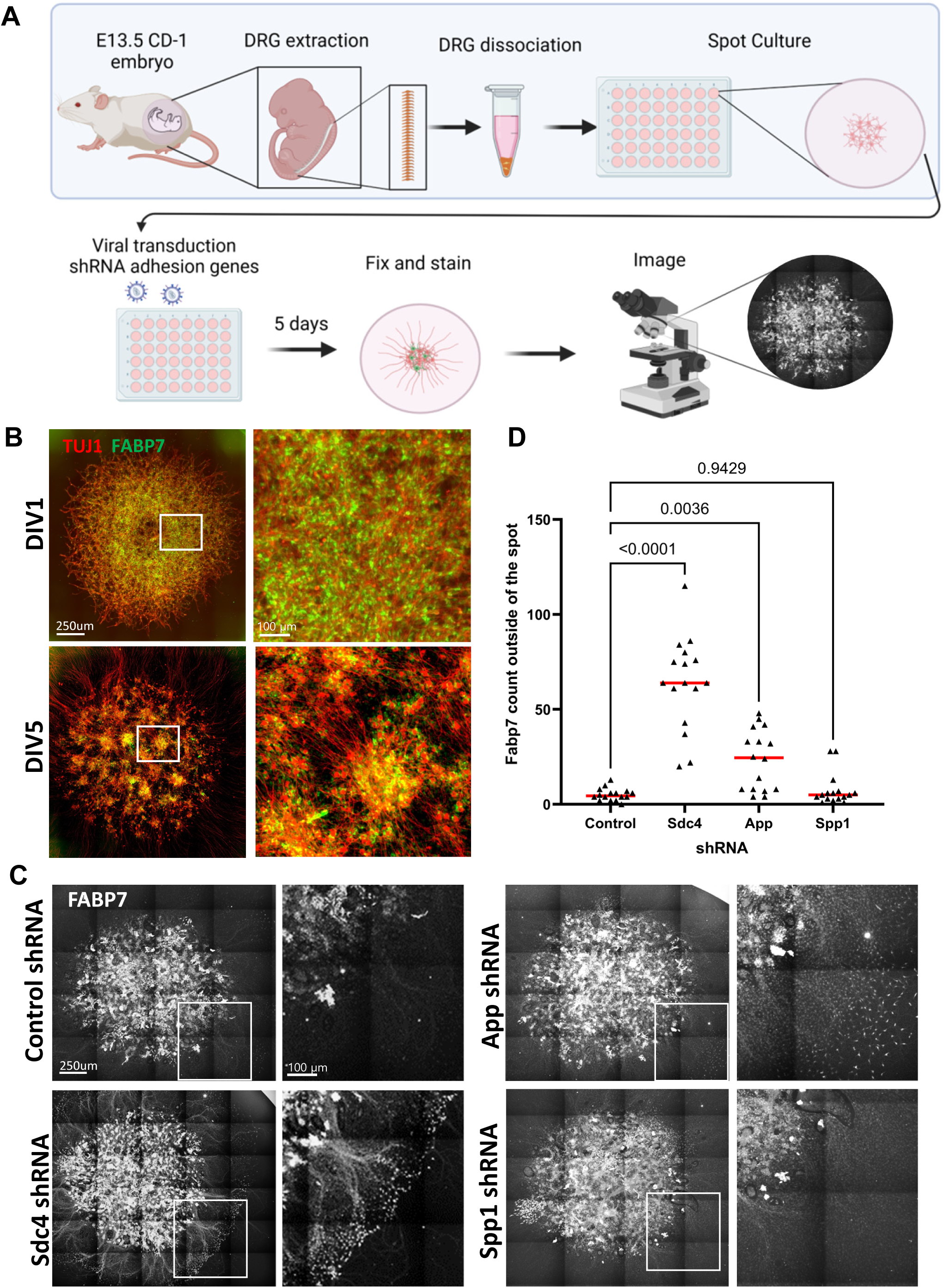
Sdc4-dependent adhesion constrains satellite glia localization in a DRG spot culture assay. **A**) Schematic of the experimental workflow for shRNA-mediated gene knockdown in embryonic DRG (eDRG) spot co-cultures, including viral transduction and imaging. **B**) Representative images of eDRG spot cultures stained for the neuronal marker TUJ1 (red) and the SGC marker FABP7 (green) at DIV1 and DIV5. Scale bars: 250 µm (low magnification) and 100 µm (high magnification). **C**) Representative DIV5 images of eDRG cultures transduced with control shRNA or Sdc4 shRNA and stained for FABP7. Scale bars: 250 µm (low magnification) and 100 µm (high magnification). **D**) Quantification of FABP7 signal outside the spot region at DIV5 across experimental conditions. n = 4 biologically independent animals. P-values were determined by one-way ANOVA.

## Discussion

Painful neuromas represent a maladaptive outcome of peripheral nerve injury, characterized by disorganized axonal regrowth, excess collagen deposition, and persistent sensory dysfunction. While neuroma pathology is commonly described at the injury site, less is known about how different post-injury interventions reshape cellular states and communication within the DRG, where sensory neuron somata and their supporting glia integrate injury signals and influence long-term pain outcomes. Here, we used single-cell transcriptomics to define the DRG response two weeks after sciatic nerve transection in a rat neuroma model and to compare two clinically relevant secondary interventions- proximal crush and neuroma resection. Across major DRG cell populations, we identify a conserved injury-associated transcriptional core coupled to intervention-specific rewiring of glial programs, immune composition, and inferred cell-cell signaling. A central theme emerging from these data is that metabolic remodeling and ECM/adhesion-mediated communication networks are major features of the DRG response. However, the relative balance between trophic and metabolic programs, which are more prominent after resection, and inflammatory programs, which are more pronounced after crush, differs markedly across injury paradigms.

### A neuroma-forming injury engages a broad DRG remodeling program

By two weeks post-transection, the injured nerve stump exhibited increased interfascicular collagen deposition and disrupted axonal organization which are hallmarks of neuroma formation, resembling changes described in human neuromas. This time point captures a phase when acute Wallerian degeneration has transitioned toward ongoing remodeling and consolidation of chronic injury-associated states. Our scRNA-seq dataset (over 63,000 cells across conditions) resolved canonical DRG populations and revealed a consistent compositional shift after injury with changes in cell-type abundance. These results align with prior studies showing that DRG injury responses involve not only neuron-intrinsic transcriptional changes but also significant remodeling of the local immune and glial milieu (Avraham et al. 2020; Avraham et al. 2021; Feng et al. 2023; Jager et al. 2020). Importantly, because cell-type proportions can be influenced by both biology and dissociation/capture biases, future validation using histology or flow cytometry will be essential to confirm absolute changes in cell number and localization.

### SGC programs diverge sharply depending on the post-injury intervention

One of the most prominent intervention-dependent effects occurred in SGCs. Following nerve cut, SGCs mounted a robust transcriptional response dominated by metabolic pathways (fatty acid/cholesterol-related programs) and glial differentiation signatures, with additional enrichment of adhesion/junction-related pathways. This pattern is consistent with the idea that SGCs respond to injured neurons by increasing metabolic support and reorganizing their ensheathment and coupling (Avraham et al. 2020; Huang, Gu, and Chen 2013; Hanani et al. 2002). In contrast, proximal crush performed two weeks after transection produced a comparatively small transcriptional response in SGCs but disproportionately enriched chemokine/cytokine and immune signaling pathways, suggesting a shift toward an inflammatory state. Neuroma resection, by comparison, largely reinforced the metabolic/differentiation programs seen after cut and showed substantial overlap with the cut response, indicating that resection engages an SGC response more similar to the primary transection injury than does crush.

This divergence has several potential interpretations. Proximal crush may introduce an additional wave of axonal degeneration/regeneration signals that preferentially recruit or activate immune-like SGC states (or immune-associated signaling within SGCs), potentially reflecting an attempt to coordinate repair and recruit myeloid cells. Resection may instead reset or reshape the local injury environment in a way that favors metabolic support and glial differentiation within the ganglion. These alternative programs are likely to have distinct consequences for pain as inflammatory signaling in SGCs can modulate neuronal excitability and may contribute to nociceptive sensitization (Blum et al. 2014), whereas metabolic and differentiation programs may be more supportive or stabilizing. Direct behavioral correlation and functional perturbation will be required to test whether the SGC state induced by crush versus resection predicts divergent pain trajectories.

### DRG-associated Schwann cells engage a conserved cholesterol biosynthesis program

Although SCs are primarily peripheral-nerve residents, DRG dissections capture DRG-associated SCs ensheathing proximal axon segments. Across all injury paradigms, this Schwann cell population upregulated a strong and conserved lipid/steroid signature, with pronounced induction of cholesterol biosynthesis pathways. This finding is consistent with the essential requirement for cholesterol and lipid metabolism during remyelination and repair (Xu et al. 2026; Saher et al. 2005), and supports the idea that even DRG-adjacent Schwann cells enact a stereotyped metabolic program after injury. Notably, canonical Schwann cell differentiation/myelination markers exhibited intervention-specific regulation, suggesting that while cholesterol biosynthesis is a shared response, the differentiation state of SCs differs by intervention. These differences may reflect how each injury strategy modulates the balance between repair Schwann cell states and remyelinating trajectories.

### Neurons activate a conserved injury program emphasizing structural remodeling and reduced excitability signatures

Despite known technical challenges in capturing neurons via whole-cell microfluidics, the neuronal dataset revealed robust differential expression across injury paradigms and, importantly, a substantial shared core of upregulated genes common to cut, crush, and resection injuries. This conserved neuronal response across all injury conditions, particularly in contrast to the more variable expression patterns observed in non-neuronal cells, suggests that environmental cues may play a significant role in regulating pain. Common pathway enrichment pointed strongly toward cytoskeletal remodeling, migration/structural organization, and adhesion/junctional programs, consistent with neurons reorganizing their somatic and axonal architecture in response to injury. The observed induction of genes associated with tight junctions and barrier maintenance (Tjp1/Tjp2 and Jam family members) raises the possibility that DRG injury provokes remodeling of the blood-nerve barrier or related microvascular interfaces within the ganglion. While DRGs are not protected by the classic blood-brain barrier, they possess specialized vascular and perineurial features (Weerasuriya and Mizisin 2011). The shared upregulation of tight junction-associated genes in neurons suggests that injured neurons may actively remodel membrane domains to form localized diffusion barriers (Förster 2008). This response may help regulate the neuronal microenvironment and control exposure to inflammatory signals that contribute to pain signaling. Our data support a model in which neuronal injury signals may participate in reorganizing local barrier-like properties as part of the response to tissue damage. In parallel, downregulated neuronal genes were enriched for ion transport pathways, including sodium, potassium, and calcium channel modules, consistent with a transient reduction in excitability-associated transcriptional programs as neurons prioritize repair and remodeling (Renthal et al. 2020).

Together, these neuronal features support a model in which the injured DRG enters a state characterized by structural reorganization and altered electrophysiological set points. Whether these transcriptional changes correspond to altered firing thresholds, spontaneous activity, or subtype-specific changes in nociceptors versus proprioceptors will be an important direction for future work, particularly given the relevance of ectopic firing and sensitization in neuroma-associated pain (North, Lazaro, and Dougherty 2018).

### Macrophages expand and reprogram metabolically, with intervention-specific features

Macrophages expanded substantially after injury, consistent with both infiltration and local proliferation (Feng et al. 2023; Kwon et al. 2013; Niemi et al. 2013), and exhibited a shared upregulation of oxidative metabolism programs (fatty acid oxidation and oxidative phosphorylation). This metabolic reprogramming could reflect macrophage adaptation to sustained tissue remodeling, debris handling, and trophic signaling within the ganglion. Interestingly, the downregulated pathways across injuries included inflammatory response signatures (cytokine/chemokine signaling, TNF, IL-17-associated modules), suggesting that at the two-week time point and later, macrophages may be transitioning away from an early inflammatory phase toward more reparative states, or that intervention strategies selectively enrich macrophage subsets with distinct inflammatory profiles. Resection uniquely enriched gene sets annotated to neuronal development, implying a macrophage state potentially tuned toward neural remodeling, while proximal crush showed reduced macrophage proliferation signatures and a smaller macrophage fraction than a cut injury. These distinctions point to intervention-dependent immune remodeling in the DRG that could influence both regeneration and pain persistence.

### Secondary interventions reshape cell-type-specific transcriptional programs in the DRG

A key insight from our study emerges from directly comparing secondary interventions to the neuroma-forming cut injury, which reveals that proximal crush and resection do not simply reverse the injury response but instead impose distinct layers of transcriptional remodeling across cell types. In SGCs, both interventions induce only modest additional changes relative to cut, but diverge in direction, with crush promoting inflammatory signaling and resection favoring regenerative and differentiation-associated programs. In contrast, Schwann cells and neurons exhibit more extensive reprogramming, with both interventions converging on pathways related to cytoskeletal organization and ECM remodeling, while diverging in inflammatory regulation. Notably, crush selectively maintains or enhances inflammatory signaling in glial compartments, whereas resection more consistently suppresses these pathways and promotes structural and trophic programs. Macrophages also show intervention-dependent tuning, with both conditions shifting toward regulated inflammatory and angiogenesis-related states compared to cut injury. Together, these findings suggest that secondary interventions reshape the DRG environment not by reverting to a pre-injury state, but by redirecting the balance between inflammatory, metabolic, and structural programs in a cell type-specific manner. This layered remodeling may be critical for determining whether the system progresses toward persistent pain or regenerative resolution, and highlights the importance of targeting cell-type-specific pathways when designing therapeutic strategies for neuroma-associated pain.

### DRG communication networks are rewired by intervention and dominated by ECM-adhesion signaling hubs

A major insight from our ligand-receptor network analysis is that the “wiring diagram” of DRG communication shifts depending on intervention. After nerve cut, upregulated interactions were concentrated within and between neurons and SGCs. In contrast, proximal crush shifted the landscape toward neuron-macrophage and neuron-Schwann cell interactions, whereas resection prominently featured immune and Schwann cell interactions alongside renewed neuron-SGC coupling. Despite these differences, a shared set of ligands and receptors was induced across all paradigms, suggesting a conserved injury communication module onto which intervention-specific programs are layered.

Across neuron-SGC interactions, ECM ligands and adhesion receptors emerged as central hubs: collagens, laminins, and thrombospondins paired with integrins and syndecans (Itgb1 and Sdc4). This is biologically plausible given the intimate perisomatic neuron-SGC association, and positions ECM-receptor signaling as a key axis linking structural remodeling to cellular signaling. The observation that resection uniquely induced pathways related to direct cell-cell communication and junction organization suggests that resection may promote a stronger contact-dependent reassembly program, potentially relevant to regeneration or stabilization of tissue architecture.

### Functional perturbation supports a causal role for adhesion genes in neuron-SGC organization

A critical step beyond inference is functional validation. Using a DRG “spot” co-culture model, we tested whether perturbing candidate adhesion/communication genes alters neuron-SGC spatial organization. Sdc4, an adhesion-related gene, was upregulated in both neurons and SGCs after cut injury, but in crush and resection paradigms its induction was restricted to neurons. Knockdown of Sdc4 in the neuron-SGC co-culture produced a clear phenotype in which SGCs redistributed beyond the spot boundary, consistent with impaired confinement and/or reduced adhesion to neuronal clusters. App knockdown showed a similar effect, whereas Spp1 knockdown did not significantly alter this spatial readout under our conditions. These results support the notion that Sdc4-dependent adhesion is not merely correlated with injury states but can shape neuron-SGC organization in a manner consistent with the signaling networks inferred from scRNA-seq.

Importantly, Sdc4 has multiple plausible ligands, both in glia and neuron compartments, including collagens and thrombospondins, several of which are induced across DRG cell types after injury (Thbs4 broadly, Thbs2 more restricted). This provides a mechanistic foothold for a model in which injury-induced ECM remodeling generates a ligand landscape that engages Sdc4/integrin receptor hubs to reorganize glial ensheathment, alter perisomatic signaling, and potentially modulate neuronal excitability (Woods and Couchman 2001; Bass et al. 2007; Lin et al. 2015). Future experiments could directly test this axis using receptor blockade, conditional knockdown/knockout strategies, or ligand- specific perturbations, combined with in vivo measurements of pain behavior and regeneration.

### Implications for neuroma biology and therapeutic intervention

Clinically, nerve crush and neuroma resection are used in different contexts and are thought to influence outcomes via distinct mechanisms. Our data suggest that these interventions also differentially tune DRG cellular states and communication networks. Proximal crush appears to bias SGCs toward inflammatory signaling and shifts interaction networks toward immune and Schwann cell engagement, potentially representing an immune-coordinated repair attempt that may also carry risk for persistent sensitization. Resection, in contrast, preferentially reinforces metabolic and differentiation programs in glia and engages junction/contact modules, which may reflect a reorganization program that could stabilize tissue architecture and support regeneration-associated communication. These distinctions generate testable hypotheses linking intervention-induced DRG states to behavioral outcomes, and point to ECM/adhesion hubs as promising therapeutic entry points.

### Limitations and future directions

This study has several limitations. First, ligand-receptor analyses infer potential interactions based on transcript abundance and curated databases, and do not directly measure protein localization, ligand availability, or receptor activation. Second, all analyses were performed at a single time point, thus temporal profiling will be essential to distinguish early inflammatory waves from later reparative programs and to identify the sequence of communication rewiring. Third, whole-cell scRNA-seq under-captures neurons. Future work using single-nucleus RNA-seq or neuron-enriched preparations will improve subtype resolution and reduce sampling bias. Fourth, our functional co-culture perturbations were performed in an embryonic mouse model, while the scRNA-seq data are from adult rat DRGs. Although the assay provides causal insight into adhesion mechanisms, species and developmental differences warrant careful interpretation and motivate in vivo validation in the relevant adult injury context.

Key next steps would include linking intervention-specific DRG states to pain behaviors and regeneration metrics, validating ECM/adhesion hubs at the protein level with spatial methods (RNAscope, immunostaining or spatial transcriptomics). Future studies should test whether manipulating Sdc4/integrin signaling alters neuron-SGC coupling, macrophage states, and long-term outcomes.

## Conclusion

In summary, our single-cell atlas of the DRG in a neuroma-forming injury model reveals that post-injury interventions are associated with distinct transcriptional and communication programs within the ganglion. Across cell types, a conserved injury response is evident, but proximal crush and neuroma resection drive divergent glial states and rewire intercellular signaling networks. ECM-adhesion signaling hubs, particularly syndecan and integrin-centered networks, emerge as dominant features of neuron-SGC communication, and functional perturbation of Sdc4 supports a causal role for adhesion genes in organizing neuron-glia interactions. These findings provide a framework for understanding how intervention strategies may shape DRG remodeling and suggest mechanistic targets for improving recovery and reducing neuroma-associated pain.

---

**Supplementary Figure 1. A**) Venn diagram of up and downregulated genes in SGCs crush and resect vs. cut injury. **B**) Pathway enrichment analysis of SGCs DEGs in crush and resect vs, cut injury. **C**) Venn diagram of up and downregulated genes in SCs crush and resect vs. cut injury **D**) Pathway enrichment analysis of SCs DEGs in crush and resect vs, cut injury.

**Supplementary Figure 2. A**) Venn diagram of up and downregulated genes in Neurons crush and resect vs. cut injury. **B**) Pathway enrichment analysis of Neurons DEGs in crush and resect vs, cut injury. **C**) Venn diagram of up and downregulated genes in Macrophages crush and resect vs. cut injury **D**) Pathway enrichment analysis of Macrophages DEGs in crush and resect vs, cut injury.

**Supplementary Figure 3.** Heatmap showing Sdc4 expression and the expression of candidate Sdc4 ligands across DRG cell types and injury paradigms.

**Supplementary table 1**- Differentially expressed genes of SGC, SCs, Macrophages and Neuron clusters in cut, crush and resect treatments compared to uninjured control and crush and resect compared to cut injury.

**Supplementary table 2-** Ligand-receptor interaction between SGC, SCs, Macrophages and Neuron clusters in cut, crush and resect treatments

**Supplementary table 3-** Ligand-receptor interaction between SGC and Neuron clusters in cut, crush and resect treatments

**Supplementary table 4-** Quantification of FABP7+ cells outside of the spot with shRNA induction

## Material and Methods

### Animals

Adult male Lewis rats (200-250 g; 7-9 weeks; Charles River Laboratory) were randomized to one of four groups (n=4). Surgical procedures and perioperative care were performed in accordance with the Institutional Animal Studies Committee and National Institutes of Health Guidelines, and all animal protocols were approved by the Washington University Animal Studies Committee (IACUC) under protocol A3381-01. All experiments were performed in accordance with the relevant guidelines and regulations. Animals were housed and cared for in the Washington University School of Medicine animal care facility. This facility is accredited by the Association for Assessment & Accreditation of Laboratory Animal Care (AALAC) and conforms to the PHS guidelines for Animal Care. Accreditation - 7/18/97, USDA Accreditation: Registration # 43-R-008. Animals were fed Rodent Diet 20 (Purina Mills Nutrition International) and water ad libitum. Animals were housed in a central climate-controlled animal care facility with a 14/10-hour light/dark cycle. They were monitored daily for signs of infection or distress by trained veterinary staff.

### Sciatic Transection/ Neuroma Formation

All surgical procedures were performed using aseptic techniques under an operating microscope. Animals were anesthetized via intraperitoneal injection of a 0.25 mL of 0.75 mg/kg ketamine and 0.5 mg/kg dexmedetomidine. In all animals on day 0, a 4 cm transverse incision was made in the mid-thigh and the sciatic nerve was exposed using a muscle-sparing approach. The sciatic nerve and its branches were mobilized and sharply transected just proximal to the trifurcation. The distal nerve was swept distally and secured to the adjacent muscle using 6-0 silk suture to decrease end-organ neurotropism. The muscle and skin were then closed in 2 layers with interrupted sutures using 4-0 Vicryl and 4-0 Nylon respectively. Rats were resuscitated following surgery with 0.1 mL intraperitoneal Antisedan and 0.1 mL of 1 mg/kg solution of buprenorphine sustained release. Rats were monitored at least daily for 7 days by lab staff for post-operative complications.

### Histomorphometry

Nerve segments for the sciatic transection group versus negative control (no sciatic injury) were fixed in an immersion of cold 3% glutaraldehyde solution in 0.1 M phosphate buffer (pH 7.2). The tissues were post- fixed with 1% osmium tetroxide, ethanol dehydrated and embedded in Araldite 502. 1µm thin sections were made from the tissue and stained with 1% toluidine blue for examination under light microscopy (Becker et al. 2000; Doolabh and Mackinnon 1999; Lee et al. 2000; Midha, Mackinnon, and Becker 1994). Slides were evaluated by an observer blinded to the experimental groups for overall nerve architecture. Cross sections from the nerve were evaluated.

### Proximal Nerve Secondary Interventions

2 weeks after sciatic nerve transection, Lewis rats were re-anesthetized, and the sciatic nerve was re-exposed using the above technique. Animals either underwent a proximal crush injury, or a resection of the developing neuroma bulb. For proximal crush injury, a #5 jeweler’s forceps was used approximately 1.5 cm proximal to the nerve transection for 30 seconds. The wound was then closed, and the animal resuscitated. For resection, the distal 3-5 mm of tissue was excised, removing the entirety of the visible neuroma. The wound was then closed, and the animals were resuscitated as above.

### Rat DRG dissociation

At 6 weeks after initial sciatic nerve transection, L4 and L5 DRGs (2 biologically independent samples, n=2 rats for each sample) were collected into cold Hank’s balanced salt solution (HBSS) with 5% Hepes, then transferred to warm Papain solution and incubated for 20 min in 37 °C. DRGs were washed in HBSS and incubated with Collagenase for 20 min in 37 °C. Ganglia were then mechanically dissociated to a single cell suspension by triturating in culture medium (Neurobasal medium), with Glutamax, PenStrep and B-27. Cells were washed in HBSS + Hepes +0.1%BSA solution, passed through a 70-micron cell strainer. Hoechst dye was added to distinguish live cells from debris and cells were FACS sorted using MoFlo HTS with Cyclone (Beckman Coulter, Indianapolis, IN). Sorted cells were washed in HBSS + Hepes+0.1%BSA solution and manually counted using a hemocytometer. Solution was adjusted to a concentration of 500 cells/µL and loaded on the 10X Chromium system.

### Single cell RNA-seq

Single-cell RNA-seq libraries were prepared using GemCode Single-Cell 3′ Gel Bead and Library Kit (10x Genomics). A digital expression matrix was obtained using 10X’s CellRanger pipeline (Build version 3.1.0) (Washington University Genome Technology Access Center). Quantification and statistical analysis were done with Partek Flow package (Build version 9.0.20.0417). Filtering criteria: Low quality cells and potential doublets were filtered out from analysis using the following parameters: total reads per cell: 600-15000, expressed genes per cell: 500-4000, mitochondrial reads <10%. A noise reduction was applied to remove low expressing genes < = 1 count. Counts were normalized and presented in logarithmic scale in CPM (count per million) approach. We applied variance stabilizing transformation to count data using a regularized Negative Binomial regression model (Seurat::SCTransform) followed by a removal of unwanted variation caused by known nuisance and/or batch factors (Scale expression). Principal component analysis (PCA) using Louvain clustering algorithm was then undertaken followed by an unbiased clustering (Graph-based clustering) algorithm implemented in Partek. Clustering was performed using Compute biomarkers algorithm, which computes the genes that are expressed highly when comparing each cluster. Seurat3 integration was used to obtain cell type markers that are conserved across samples and clusters were assigned to a cell population by at least three established marker genes. Clusters are presented in t-SNE (t-distributed stochastic neighbor embedding) plot, using a dimensional reduction algorithm that shows groups of similar cells as clusters on a scatter plot. Differential gene expression analysis performed using three different models: Compute biomarkers following regularized Negative Binomial regression, non-parametric ANOVA and the Partek algorithm GSA that integrate multiple statistical models. Gene lists from each statistical model were intersected to remove potentially false positive genes. The intersected lists were then applied for all downstream analyses. A gene was considered differentially expressed (DE) if it has a false discovery rate (FDR) step-up (p-value adjusted). p ≤ 0.05 and a Log2 fold change ≥ ± 2. The DE genes were subsequently analyzed for enrichment of GO terms and the KEGG pathways using Partek flow pathway analysis. Partek was also used to generate figures for t-SNE and scatter plot representing gene expression. GO and KEGG tools were used for pathway analysis for DEG and SRplot was used to generate the bar and heatmap plots(Tang et al. 2023).

### Embryonic mouse DRG spot cultures

Dorsal root ganglia were isolated from time pregnant e13.5 CD-1 mice (Charles River Laboratories) into dissection media consisting of DMEM and Pen/Strep. After a short centrifugation, dissection media was aspirated and cells were digested in 0.05% Trypsin-EDTA for 25 min. Next, cells were pelleted by centrifuging for 2 min at 500 × *g*, the supernatant was aspirated, and Neurobasal was added. Cells were then triturated 25x and added to the growth medium containing Neurobasal, B27 Plus, 1 ng/ml NGF, Glutamax, and Pen/Strep. Approximately 10,000 cells were added to each well in a 2.5 μl spot. Spotted cells were allowed to adhere for 10 min before the addition of the growth medium. Plates were pre-coated with 100 μg/ml poly-D-lysine overnight and washed with sterile water prior to plating. Viral particles containing shRNA were added to the media at DIV1. For immunostaining, wells were incubated in PBS-0.1% Triton (PBST) for 1 h at room temperature containing primary antibody. The wells were then washed 3x with PBST and then incubated in PBST solution containing fluorescent-labeled goat anti-rabbit secondary antibody for 1 h at room temperature. Finally, the wells were washed 3x with PBS. For immunostaining of spot culture at DIV1 and DIV5, wells were stained with TUJ1 primary antibody (Biolegend 802001; RRID:AB_2564645) (1:1,000), Fabp7 (Thermo Fisher Scientific Cat# PA5-24949, RRID:AB_2542449) (1:1,000) and DAPI. shRNA assay performed with Mission shRNA (Sigma) lentiviral transduction particles containing shRNA against mouse genes and non-target shRNA.

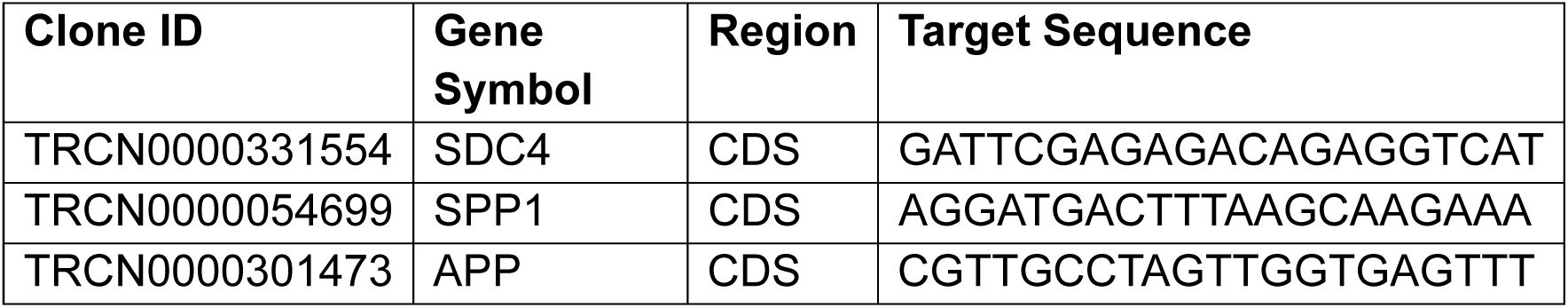

Images were acquired using an EVOS™ M7000. Cropped images of 300×300 pixels were uploaded into ImageJ software to quantify fluorescence. The analysis parameters were set to measure pixels with 10 as the lower boundary and infinity as the upper boundary, and normalized to background.

## Data availability

The raw Fastq files and the processed filtered count matrix for scRNA sequencing were deposited at the NCBI GEO database under the accession number GSE169301, GSE315353.

## Acknowledgements

This work was supported in part by the Franklin College of Arts and Sciences and the Department of Cellular Biology at the University of Georgia to OA, and by a Pilot Project Award from the Hope Center for Neurological Disorders at Washington University to VC.

## Author Contributions

Conceptualization: AM, AH, OA, VC; Data generation and analysis: AH, CS, MM, OA; Funding acquisition: VC, AM, OA; Supervision: VC, AM, OA; Writing: AM, VC, OA

