## Supplementary figures and images for "Single-Cell Atlas of Dorsal Root Ganglion Remodeling After Neuroma-Forming Nerve Injury Reveals Intervention-Specific Glial and Immune Programs"

### Supplemental Figures

#### Supplement figure1

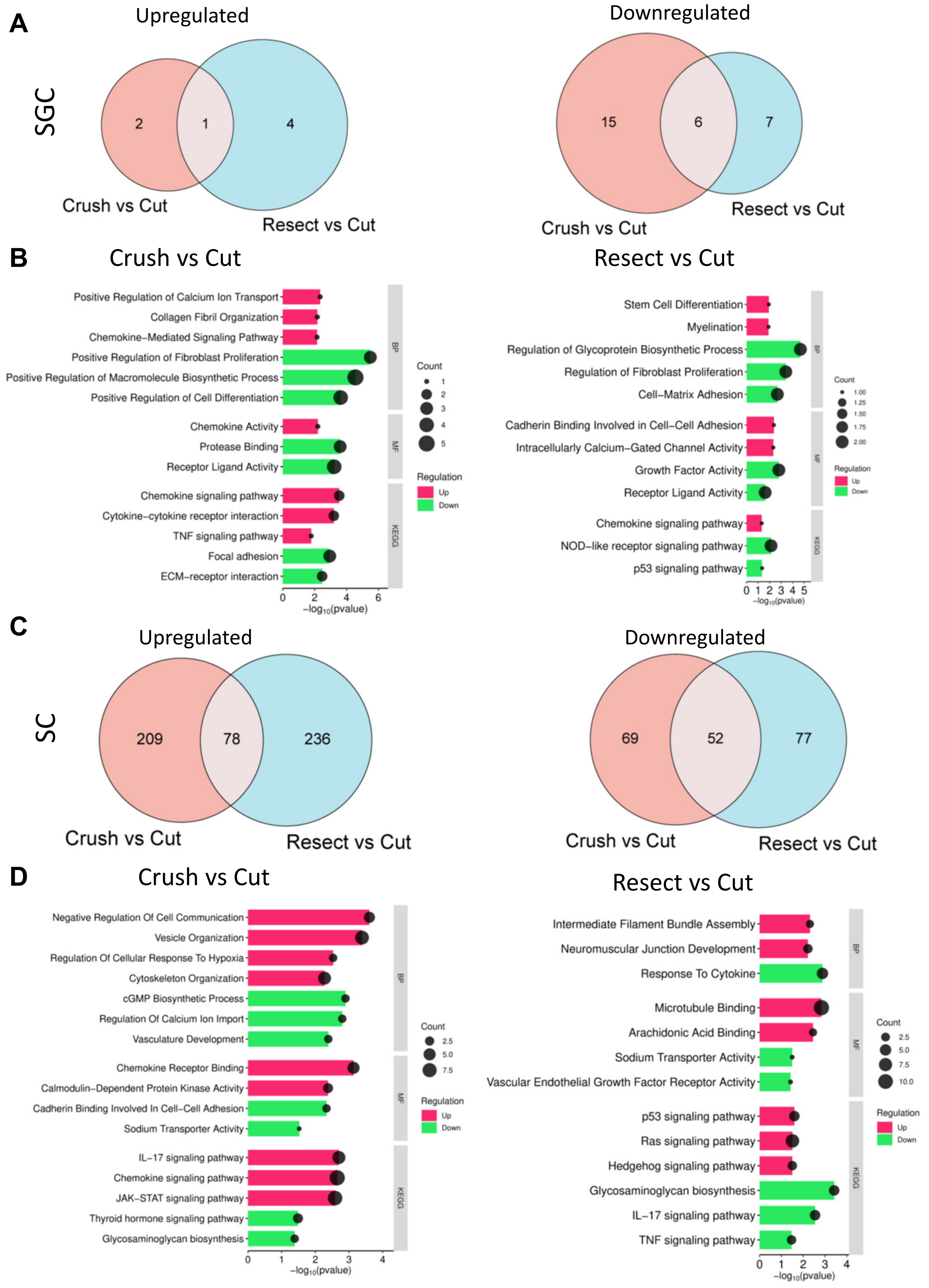

#### Supplement figure 2

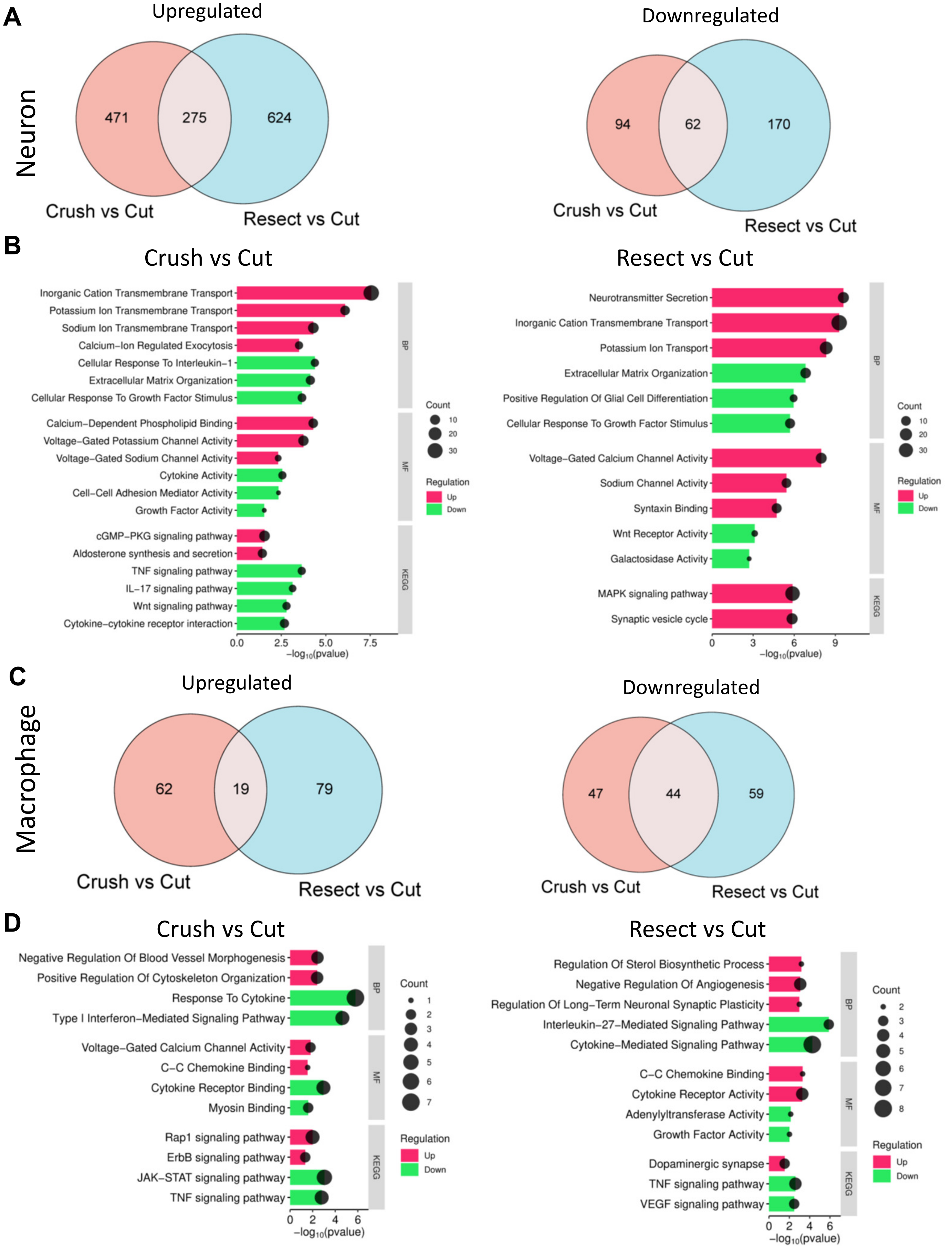

**Supplement figure 3**

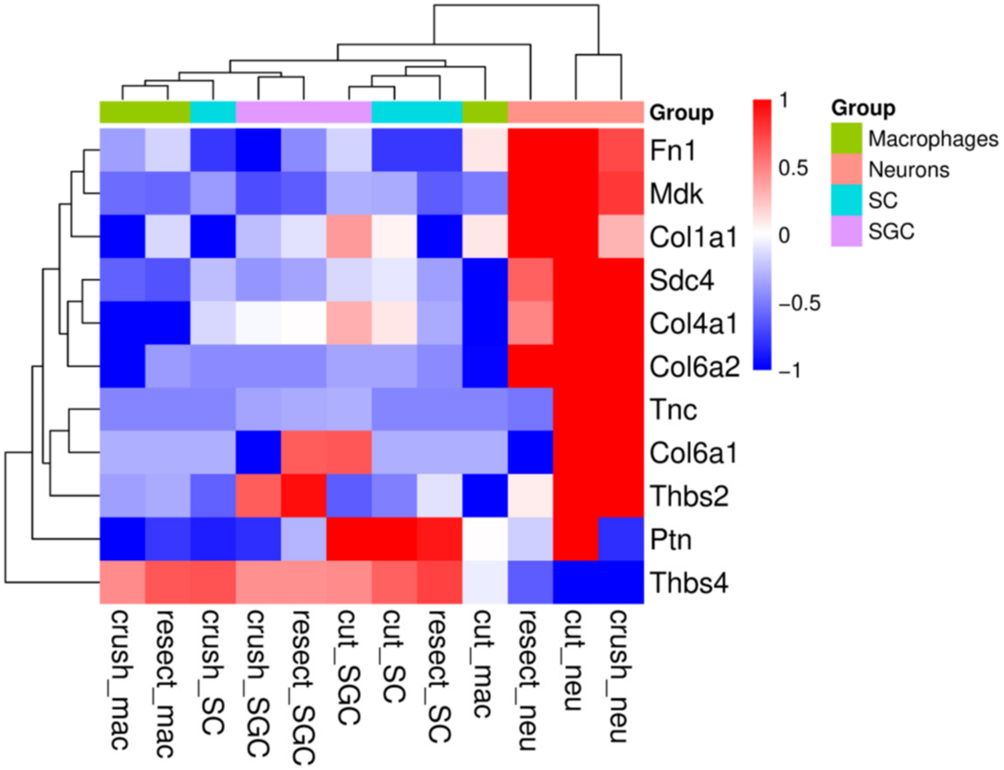
